# Targeting and Imaging Mitochondria with a Novel Cyanine-based Fluorescent Dye and Its Application to Whole Brain Imaging

**DOI:** 10.1101/2023.09.01.555877

**Authors:** Xin Yan, Xinqian Chen, Zhiying Shan, Lanrong Bi

## Abstract

In vivo, brain imaging poses several challenges due to the complexity of brain tissue and the requirements for effective and safe imaging. In this manuscript, we reported on developing a new fluorescent dye, Cy5-PEG2, that can selectively accumulate in mitochondria, allowing us to visualize these organelles in different cell lines. Its versatility makes it broadly applicable in various experimental conditions, including monitoring mitochondrial dynamics in living cells. Moreover, we demonstrated that the Cy5-PEG2 dye could penetrate the blood-brain barrier (BBB), making it a promising tool for in vivo brain imaging. We further examined the response of glial cells in the hippocampus and neocortex regions of the brain through biomarkers GFAP and Iba1 but found no significant increase in their expression levels. This indicates that Cy5-PEG2 dye had no noticeable adverse effects on the brain’s immune response or neuronal health. We anticipate such a mitochondria-targeting fluorescent dye can be used to facilitate the study of mitochondrial dynamics and function in the context of whole-brain physiology and disease progression. Further research into the safety and efficacy of this dye may be necessary.

## Introduction

Mitochondria-targeting fluorescent dyes can help scientists observe and track changes in mitochondrial health and form in real-time, providing vital information on the function and dysfunction of mitochondria in many diseases. This is especially crucial for neurodegenerative diseases, as mitochondrial dysfunction is often an initial symptom.^1,2^ Detecting even minor changes in mitochondrial function through non-invasive methods is essential for early detection and effective intervention. Creating specific fluorescent dyes for each disease can also enable personalized diagnosis and treatment approaches tailored to each patient. This will be a significant achievement if successful since neurodegenerative diseases are typically complex and varied, and personalized approaches are essential for successful treatment.

The use of fluorescent dyes in bioimaging has significantly transformed the field, particularly with the introduction of cyanine derivatives. Cyanine-based fluorescent dyes are designed to emit light when excited by a specific wavelength, making them valuable tools for biological imaging.^3^ We are currently focused on expanding the library of these derivatives to develop improved dyes that possess enhanced properties such as increased brightness and biocompatibility.

It is known that dysfunctional mitochondria are linked to neurodegenerative diseases, and specific changes in these organelles can serve as biomarkers for disease progression. ^1,2,4^ We strived to develop new mitochondria-targeting fluorescent dyes to help detect and monitor these biomarkers. Previous studies showed that the accumulation of amyloid beta (Aβ) peptide in the brains of individuals with Alzheimer’s disease (AD) significantly contributes to the disease’s development and progression.^5,6^ It is believed that Aβ accumulation leads to mitochondrial dysfunction, further exacerbating neurodegeneration. ^5,6^

This manuscript reported the synthesis and biological evaluation of our newly synthesized mitochondria-targeting fluorescent dye, Cy5-PEG2. We conducted biological evaluations of this dye and used it to monitor changes in mitochondrial morphology and dynamics under normal and Alzheimer’s disease-like conditions, specifically in the presence of the Aβ_1-42_ peptide. We also investigated the ability of the Cy5-PEG2 dye to penetrate the blood-brain barrier (BBB) and be used for in vivo brain imaging.

## Materials and Methods

### Synthesis Procedure

The experiment utilized reagents and solvents obtained from commercial sources and used as received unless otherwise specified. Thin-layer chromatography (TLC) was carried out on Sigma-Aldrich TLC plates, which were silica gel and had a plate number of 1 over the glass support, with a thickness of 0.25 μm. Flash column chromatography was performed with Alfa Aesar silica gel, which had a particle size of 230–400 mesh. Melting points were measured using a MELTEMP melting point apparatus and were uncorrected. The 1H and 13C nuclear magnetic resonance (NMR) spectra were measured using a Varian UNITY INOVA instrument at 400 and 100 MHz, respectively. The chemical shifts (δ) were reported in reference to solvent residual peaks (dimethyl sulfoxide (DMSO)-d6, 1H, δ = 2.50 ppm, 13C, δ = 39.52 ppm). The data were reported as follows: chemical shift, multiplicity (s = singlet, d = doublet, t = triplet, q = quartet, m = multiplet), coupling constant J, and integration. High-resolution mass spectra (HR-MS) were obtained on a JEOL JMS HX 110A mass spectrometer. UV–vis spectra were recorded on a PerkinElmer Lambda 35 UV/Vis Spectrometer equipped with a PTP 1+1 Peltier Temperature Programmer accessory at 37 °C. Fluorescence spectra were obtained with a Horiba Jobin Yvon Fluoromax-4 spectrofluorometer using an excitation wavelength of 360 nm and slit widths of 5 nm.

#### Synthetic procedure for N-((1E,3E)-3-(phenylimino) prop1-en-1-yl)aniline (compound 1)

To a stirring solution of distilled water (100ml), HCl(20ml), and 1,1,3,3-tetramethoxypropane (10.0g, 0.11mol), a combination of distilled water (100ml), HCl (25ml) and aniline (15g, 0.10mol) was added dropwise and continued stirring around 50 °C for 3 hours. After filtration, the filtrate was further washed with ether to yield compound **1** (8.8g, 50%) as an orange powder. ^1^H NMR (400 MHz, DMSO-d_6_) 6.53 (t, *J* = 11.5 Hz, 1H), 7.16 (t, *J* = 7.2 Hz, 2H), 7.26 – 7.62 (m, 8H), 7.16 (t, *J* = 7.2 Hz, 2H). ^13^C NMR (101 MHz, DMSO-d_6_) δ 159.00, 139.31, 130.42, 126.44, 118.04, 99.34.

#### Synthetic procedure for 2,3,3-trimethyl-3H-indole (compound 2)

To a stirring solution of phenylhydrazine (3.24 g, mol), glacial acetic acid (25 ml) was added 3-methylbutanone (5ml, 0.035mol). Then, the mixture was kept refluxed at 118°C. The mixture was subjected to evaporation using a rotary evaporator, and the resulting residue was separated using 50 ml of ether and 50 ml of water. The water layer was rewashed with ether. The combined organic layers were then evaporated until they dried out, resulting in a yield of 4.0g of compound **2** (83%), which was a brownish liquid.^1^H NMR (400 MHz, CDCl_3_) δ ppm 1.26 (s, 6H), 2.24 (s, 3H), 7.15 (t, *J* = 7.4 Hz, 1H), 7.25 (dd, *J* = 8.9, 4.1 Hz, 2H), 7.51 (d, *J* = 7.6 Hz, 1H). ^13^C NMR (101 MHz, CDCl_3_) δ 188.17, 153.56, 145.69, 127.74, 125.30, 121.45, 119.99, 119.68, 53.84, 23.37, 15.62.

#### Synthetic procedure for 1-ethyl-2,3,3-trimethyl-3H-indol1-ium (compound 3)

To synthesize 1-ethyl-2,3,3-trimethyl3H-indol-1-ium, a SN2 reaction was proceeded between 2,3,3-trimethyl-3H-indole and ethyl iodide in the presence of acetonitrile as the solvent. To create a homogeneous reaction mixture, the first step involves dissolving 2,3,3-trimethyl-3H-indole in acetonitrile. Next, the deprotonated nitrogen atom in 2,3,3-trimethyl-3H-indole acts as a nucleophile and attacks the carbon atom in ethyl iodide, forming a new carbon-nitrogen bond. The iodide ion is displaced as a leaving group, and the rearrangement of the molecule stabilizes the positive charge on the nitrogen atom.

4.0 g of compound **2** was dissolved in 50 ml of acetonitrile, and 5 ml of ethyl iodide was added. The mixture was then refluxed under nitrogen protection for 48 hours. After evaporation with a rotatory evaporator, the crude residue was washed with ether to obtain compound **3** as an orange powder, with a yield of 4.2g (89%).^1^H NMR (400 MHz, CDCl_3_) *δ* (ppm): 1.54-1.58 (m, 9H), 3.06 (s, 3H), 4.66 (m, 2H), 7.51 (m, 3H), 7.54 (m, 1H).

#### Synthetic procedure for 1-(5-carboxypentyl)-2,3,3-trimethyl-3H-indol-1-ium (compound 4)

To a stirring solution of potassium iodide (3.70g, 22 mmol) in 30 ml acetonitrile, 6bromohexanoic acid (2.0g, 10 mmol) was added dropwise and kept stirring at 50 °C. After gently heating for 0.5 hours, 2,3,3-trimethyl-3H-indole (1.6g, 10 mmol) was added, and the mixture was refluxed overnight. After cooling to room temperature, the crude residue was washed with dichloromethane and concentrated with a rotary evaporator. The obtained residue was then washed with ether to yield compound **4** (3.12g, 78%) as a gray solid.^1^H NMR (400 MHz, DMSO-d_6_) δ ppm 1.37 (d, *J* = 17.5 Hz, 2H), 1.51 (m, 6H), 1.82 (p, *J* = 7.9 Hz, 2H), 2.19 (q, *J* = 6.7, 6.1 Hz, 2H), 2.47 (td, *J* = 3.3, 2.8, 1.4 Hz, 2H), 2.82 (s, 3H), 4.43 (t, *J* = 7.7 Hz, 2H), 7.56 – 7.64 (m, 2H), 7.73 – 7.89 (m, 1H), 8.90 – 8.00 (m, 1H).

#### Synthetic procedure for compound 5

To a stirring solution of compound **3** (1g, 3.2mmol) in acetic acid and acetic anhydride, compound **1** (1g, 3.8mmol) was added and refluxed under N_2_ for 1h. After cooling to room temperature, a solution of compound **4** (2.0g, 5mmol) in anhydrous pyridine was added to the mixture and continued stirring under N_2_ at room temperature overnight. After precipitation with cold ether, the green solution was decanted, leaving the crude compound **5** (∼3.2mmol) as a green solid directly used for the next steps.

#### Synthetic procedure for compound 6

To a stirring solution of crude compound **5** (∼3.2mmol) in 50ml dichloromethane was sequentially added 2-(2-(but-3-yn-1-yloxy) ethoxy)ethan-1-amine (0.62ml, 4.3mmol), 2-(1H-Benzotriazol-1-yl)-1, 1, 3, 3-tetramethyluroniumhexafluoro-phosphate, HBTU) (2.0g 5mmol) and 5ml triethylamine. The mixture was gently stirred overnight. After evaporated solvent with rotary evaporator, the crude residue was further purified via flash chromatography on silica gel (eluent: DCM: MeOH = 40:1) to yield compound **6** as a green solid.^1^H NMR (500 MHz, CDCl3) δ ppm 1.36-1.38(m, 3H), 1.42-1.44(m, 2H), 1.64-1.66(m, 14H), 1.66-1.74(m, 2H), 2.15 (t, *J* = 7.5 Hz, 2H), 2.37 (t, *J* = 2.5 Hz, 1H), 3.31-3.34 (m, 2H), 3.44-3.46(m, 2H), 3.52-3.62(m, 4H), 3.87-4.21(m, 4H), 6.02-6.09(m, 2H) 6.35 (t, J = 2.5 Hz, 1H), 6.55(t, *J* = 15 Hz, 1H), 7.01-7.05(m, 2H), 7.09-7.15(m, 2H), 7.24-7.28(m, 4H), 7.87 (td, *J* = 4, 2.8, 1.4 Hz, 2H):^13^C NMR (101 MHz, DMSO-d_6_) δ 173.08, 172.91, 172.67, 165.65, 153.40, 153.23, 141.90, 141.51, 141.36, 141.22, 128.65, 128.63, 125.79, 125.21, 125.14, 122.29, 122.23, 110.57, 110.41, 103.20, 103.06, 79.62, 77.56, 77.31, 77.05, 74.79, 69.90, 69.52, 69.05, 58.29, 49.39, 49.33, 44.01, 39.13, 39.00, 38.55, 35.98, 27.78, 27.68, 27.00, 26.32, 25.13, 12.22.

### Cell Culture

HEK293 cells were purchased from the American Type Culture Collection (ATCC). The HEK293 cells were cultured in Dulbecco’s Modified Eagle’s Medium (DMEM) supplemented with 10% Fetal Bovine Serum (FBS) and 1% penicillin/streptomycin. Growth media were replaced every 2-3 days.

Human umbilical vein endothelial cells (HUVEC) were purchased from the ATCC. The HUVECs were then cultured in DMEM, supplemented with 5.5 mM glucose, 10% FBS, 100 U/ml penicillin, and 100 mg/ml streptomycin. They were kept at 37°C in a 5% CO_2_-humidified incubator to maintain the cells.

HeLa cells were purchased from the ATCC. To maintain them for the imaging study, RPMI-1640 medium was used, and then 10% fetal bovine serum, 2 mM glutamine, 100 μg/mL penicillin, and streptomycin were added. The cells were kept in a humid atmosphere with 5% CO_2_ at 37 °C.

### Subcellular Localization Study of Cy5-PEG2 Dye

To cultivate cells on coverslips, place them in a Petri dish filled with culture medium. Once the cells have reached the desired density, remove the medium from the dish and add a pre-warmed (37°C) staining solution that comprises a MitoTracker (1 *μ*M)/LysoTracker probe (1 *μ*M), Cy5-PEG2 dye (1 *μ*M), and Hoechst 33342 (1 *μ*g/mL). After incubating the cells for 30 minutes, replace the staining solution with fresh, pre-warmed media. Finally, examine the cells using a confocal laser scanning fluorescence microscope. To analyze the obtained images, employ the Image J analysis software. Quantify the co-localization of the fluorescent signals from the dyes using a MitoTracker/LysoTracker probe and the Cy5-PEG2 dye. This provides information about the location of the Cy5-PEG2 dye within the cells.

### Animals

Adult Sprague Dawley (SD) rats used in this study were purchased from Charles River Laboratories (Wilmington, MA). All animals were obtained on a 12:12-h light-dark cycle in a climate-controlled room, and chow and water were provided *ad libitum*. This study was carried out in strict accordance with the recommendations in the Guide for the Care and Use of Laboratory Animals of the National Institutes of Health. The protocol was approved by the Michigan Technological University Institutional Animal Care and Use Committee.

### Primary neuronal cell culture

The primary neuronal cells were isolated from the brain tissues of 1∼2-day-old SD pups using the papain dissociation system (Worthington Biochemical Corporation) as instructed by the manufacturer. These neurons were then cultured in Neurobasal-A medium supplemented with 2% B27+ and 1% antibiotic-penicillin/streptomycin on poly Dlysine-coated 35 mm dishes. The cells were incubated at 37°C in an incubator containing 5% CO2, and half medium was replaced every 3 days. After 7∼10 days, the neuronal cultures were incubated with Aβ1-42 peptide at varying concentrations for 24h and then co-stained with the Cy5PEG2 dye (1 *μ*M, green fluorescence) and Hoechst (0.1 *μ*g/mL, blue) for the study of mitochondrial morphology and dynamics.

### sIntraperitoneal (IP) injection and fluorescent immunostaining

The Cy5-PEG2 dye diluted in PBS was administered into the inguinal area of SD rats. SD Rats were restrained appropriately in the head-down position. The peritoneal cavity in the lower right quadrant of the abdomen, lateral to the rat’s midline, was the injection site. Rats were held with the body tilted downward, and the head tilted back to slide organs forward. 24 hours following Cy5-PEG-2 administration, SD rats were perfused with ice-cold PBS and 4% paraformaldehyde (PFA). The rat brains were fixated overnight with 4% PFA and subjected to 30% sucrose in filtered PBS until the tissue sank. Then, the brains were cryo-sectioned into 15-20 µm thick coronal sections and subjected to fluorescent immunostaining as described in our previous publication. In brief, sectioned cortex or hippocampus were washed in PBS and then incubated with 5% horse serum for 30 minutes. Subsequently, brain sections were incubated with either rabbit anti-NeuN antibody (Cell signaling, 1:300 dilution) or rabbit antiGFAP antibody (Abcam, 1:500 dilution), or rabbit anti-IBA1 antibody (Fujifilm Wako, 1:500 dilution) in PBS containing 0.5% Triton X-100 and 5% horse serum for 48 h at 4°C. Brain sections were rewashed with PBS and then incubated with secondary antibody Alexa fluor 488 donkey anti-rabbit IgG at room temperature for 1h. The immunoreactivity of NeuN, GFAP, IBA1, and fluorescence of Cy5-PEG-2 was observed under a confocal laser scanning microscope (Olympus).

## Results

### The rationale of molecular design

Asymmetric cyanine dyes are highly favored over symmetric ones for several reasons. ^3,7^ They produce brighter, more sensitive signals and better absorption and emission characteristics. Their longer wavelengths make them less susceptible to background interference and improve light penetration through biological samples. Furthermore, asymmetric cyanine dyes have large Stokes shifts, which reduces background noise and improves discrimination between excitation and emission light. ^3,7^ As a result, we have developed a series of new fluorescent dyes, such as Cy5-PEG2, based on the asymmetric cyanine structure.

The PEG linker is widely used for biomedical applications due to its biocompatibility and ability to improve drug delivery systems. The polyethylene glycol units in the PEG linker play a vital role in enhancing the ability of drug delivery systems to target mitochondria. Studies have shown that PEGylation can increase the hydrophilicity of a molecule, reduce non-specific interactions with cell membranes, and improve cellular uptake.^8^ To improve the targeting capacity of the asymmetric cyanine derivative, we incorporated a PEG linker to enhance cellular uptake, protect the fluorescent dye from degradation, and improve mitochondria-targeting capacity.

### Synthesis of the Cy5-PEG2 dye

**Scheme 1.**
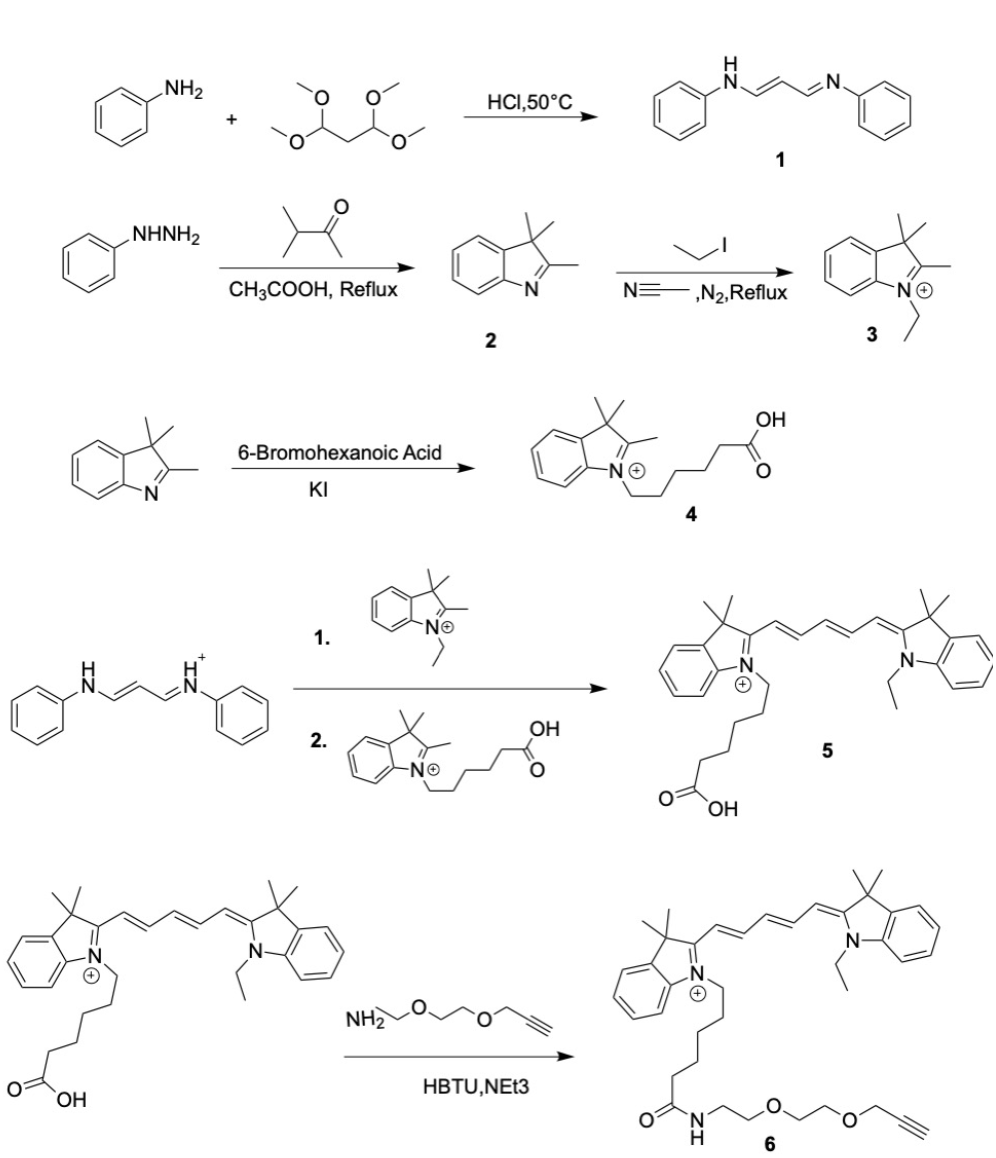
Synthetic scheme of preparation of the Cy5PEG2 dye.

Using HCl, 1,1,3,3-tetramethoxypropane (TMP), and aniline to create compound **1**, this reaction involves the formation of an iminium ion intermediate, followed by an intramolecular cyclization reaction and a rearrangement and deprotonation step. Using phenylhydrazine, glacial acetic acid, and 3-methylbutanone to prepare compound **2**, this process involves forming phenylhydrazine acetate followed by adding phenylhydrazine acetate to 3-methylbutanone to form an imine intermediate, which then undergoes protonation and rearrangement to form a more stable enamine intermediate. The enamine intermediate subsequently undergoes intramolecular cyclization, resulting in the formation of compound **2**. To create compound **3**, an SN2 reaction is necessary between compound **2** and ethyl iodide while using acetonitrile as the solvent. In this reaction, the deprotonated nitrogen atom in compound **2** acts as a nucleophile to attack the carbon atom in ethyl iodide, forming a new carbon-nitrogen bond. Then, the iodide ion is displaced as a leaving group, and the rearrangement of the molecule stabilizes the positive charge on the nitrogen atom. The synthesis of compound **4** also involves a multistep reaction mechanism. In this process, a potassium bromide salt is first formed and then reacts with an acyl iodide intermediate through a nucleophilic substitution reaction between 6-bromohexanoic acid and potassium iodide. The second step is a nucleophilic addition reaction between the acyl iodide and 2,3,3-trimethyl-3H-indole, forming a tetrahedral intermediate. The third step involves the rearrangement of the tetrahedral intermediate to a zwitterionic intermediate, followed by the elimination of a proton to form the desired product. Asymmetric cyanine dye **5** was synthesized using a known condensation reaction of the carboxyindoleninium salt **4**, compound **3**, and malonaldehyde dianil **1**. This process involves incorporating the quaternary ammonium group into the cyanine core and subsequent alkylation of the tertiary amino group. Finally, compound **5** proceeded with the coupling reaction with a PEG linker to provide the Cy5-PEG2 dye **6** (**Scheme 1**).

### Biological Evaluation

When considering a new fluorescent dye for monitoring changes in mitochondrial morphology and dynamics during cellular assays, assessing a few critical factors is essential. One crucial aspect is the dye’s ability to accumulate selectively in mitochondria while reducing its binding to other cellular components. This ensures precise labeling of mitochondria and specificity for this organelle. Additionally, the dye should penetrate cells and mitochondrial membranes without causing toxicity or interfering with mitochondrial functions. An optimal fluorescent dye for live-cell imaging should be photostable, exhibit high fluorescence brightness, and contrast against the cellular background. Lastly, evaluating the dye’s compatibility with the specific experimental conditions of the cellular assays is vital. Herein, we conducted various assays to determine whether the Cy5-PEG2 dye suits live cell imaging assays.

### Mitochondrial targeting property of Cy5-PEG2 dye

First, we examined the subcellular localization of Cy5PEG2 dye in HEK293 cells, derived from human embryonic kidney cells and commonly used in cell biology research for various applications. HEK293 cells are a popular choice because they are easy to culture and grow quickly, and their relatively large size and flat morphology make them suitable for cell imaging.

We performed a cellular imaging experiment using a confocal laser scanning fluorescent microscope and Cy5PEG2 dye at a concentration of 1 *μ*M. The dye was incubated for 30 minutes, producing bright fluorescence and fast cellular internalization (**Fig. 1**). Furthermore, the Cy5PEG2 dye showed remarkable photostability, allowing us to carry out prolonged confocal imaging without considerable photobleaching on HEK293 cells.

**Figure 1.**
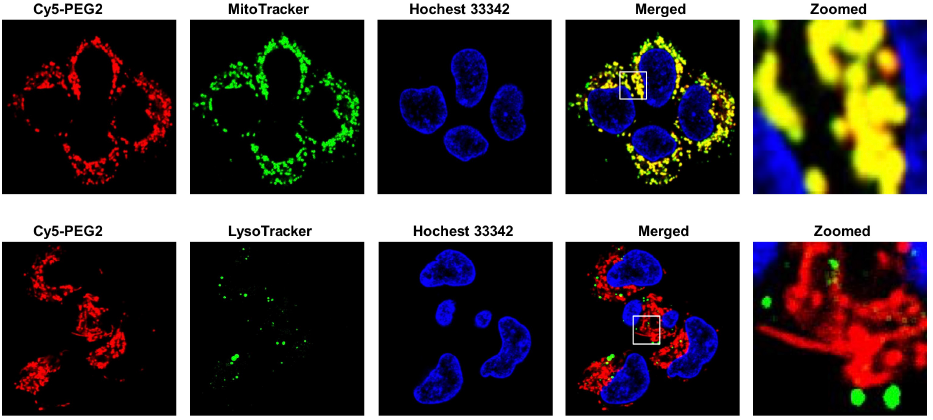
Intracellular distribution of the Cy5-PEG2 dye compared to MitoTracker and LysoTrcker. HEK293 cells incubated with the Cy5-PEG2 dye (, red fluorescence), followed by counterstaining with MitoTracker (, row 1)/LysoTracker (green fluorescence, row 2), Hoechst 33342 (1 *μ*g/mL, blue fluorescence) for 30 minutes; The images were taken at randomly selected areas to ensure the imaging conditions were consistent throughout the experiments. To ensure the reliability of the results, the experiments were repeated four times. Cells were imaged on an inverted laser scanning fluorescent microscope (Olympus) using a 60× oil immersion objective lens.

When conducting colocalization studies, we examined the spatial overlap of two molecules within a cell. Our present study looked at the colocalization of Cy5-PEG2 dye and MitoTracker, a probe commonly used to label mitochondria. We acquired fluorescent images of the samples using a confocal laser scanning microscope system. To ensure the accuracy of our results, we used ImageJ image analysis software to analyze the acquired images. Our analysis allowed us to determine the extent to which the two fluorescent signals overlapped. We used Person’s colocalization coefficient (PCC) to quantify this overlap, which calculates coefficients or pixel intensities that indicate the degree of overlap between the two signals. The fluorescence image produced using Cy5-PEG2 dye exhibits high levels of colocalization with that of MitoTracker [Person’s correlation coefficient (PCC_HEK293_): 0.90 in HEK 293 cells]. The observed high Person’s correlation coefficient established the mitochondrial specificity of the two probes. To further confirm the organelle-specificity of Cy5-PEG2 dye, we stained the cells with Lysotracker, which labels explicitly lysosomes. Our observation that Cy5-PEG2 dye overlaps with MitoTracker but not with Lysotracker suggests that the Cy5-PEG2 dye specifically stains mitochondria and not lysosomes (**Fig. 1**).

To further examine the fluorescent dyes’ subcellular localization, it is beneficial to use different cell lines. By examining the performance of the Cy5-PEG2 dye in different cell types, we can determine whether the observed localization is consistent across various cellular environments or specific to a particular cell line. We compared the localization patterns across different cell types (HUVEC, HEK293, and HeLa cell lines) and noted the similarities. We further performed statistical analysis to quantify and compare the localization patterns quantitatively. This analysis involved calculating colocalization coefficients (PCC_HUVEC_: 0.88 and PCC_HeLa_: 0.92, Supporting Information) and measuring signal intensity. We finally confirmed the subcellular localization of Cy5-PEG2 dye is consistent across the tested cell types (HUVEC, HEK293, and HeLa cell lines). We then repeated the experiments using multiple replicates to ensure the reliability of our results. These results could provide solid evidence to support the subcellular localization pattern of the Cy5-PEG2 dye observed.

By observing changes in the distribution of the Cy5PEG2 dye within cells over time, we can capture the dynamic processes within cells. This method helps us validate our initial observations by examining whether the observed localizations remain consistent or change over time. We treated HEK293 cells with the Cy5-PEG2 dye at the predetermined concentration (1 *μ*M) and incubation time (1 h, 12 h, 24 h, and 48 h). We then imaged the cells using a confocal fluorescence microscope at each time point (**Fig. 2 & 3**). We captured images of randomly selected areas. We ensured consistent imaging conditions throughout the experiment. We analyzed the acquired images using ImageJ analysis software and found no significant changes in fluorescent intensity and subcellular localization after incubation with the Cy5-PEG2 dye for an extended time. This indicates that this dye did not affect mitochondrial function or cellular processes over time.

**Figure 2.**
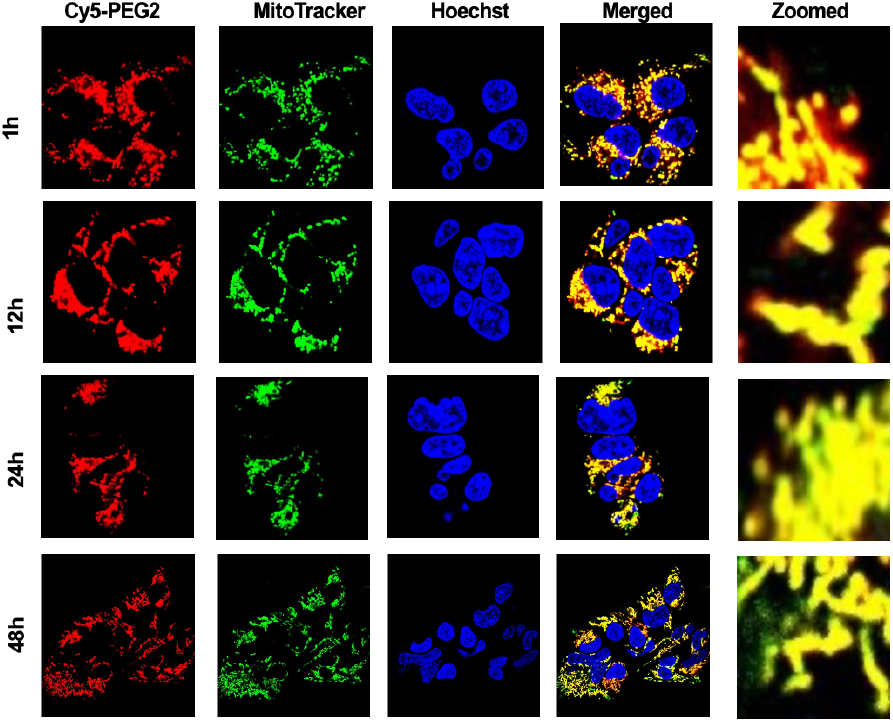
Representative confocal images of the HEK293 cells were taken at different time points (1h, 12h, 24h, and 48h). The cells underwent a time-course study and were stained with the Cy5-PEG2 dye (1 *μ*M, red fluorescence) and co-stained with MitoTacker (1 *μ*M, green fluorescence) and Hoechst (1 μg/mL, blue fluorescence). The images were taken at randomly selected areas to ensure the imaging conditions were consistent throughout the experiments. To ensure the reliability of the results, the experiments were repeated four times. Cells were imaged on an inverted laser scanning fluorescent microscope (Olympus) using a 60× oil immersion objective lens.

**Figure 3.**
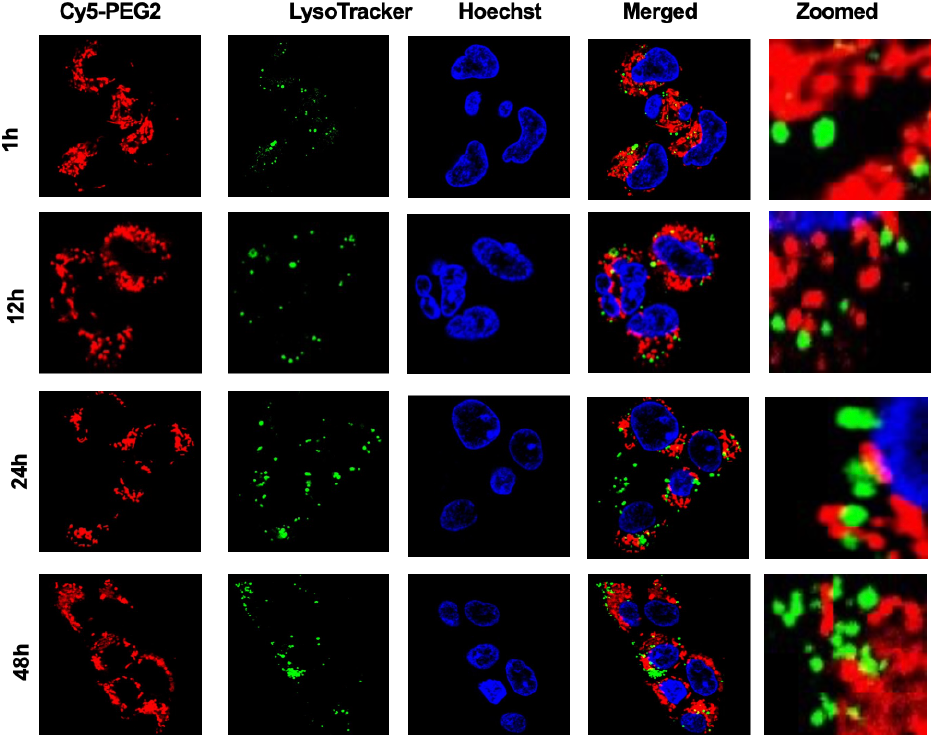
Representative confocal images of the HEK293 cells were taken at different time points (1h, 12h, 24h, and 48h). The cells underwent a time-course study and were stained with Cy5-PEG2 (1 μM, red fluorescence) and costained with LysoTacker (1 μM, green fluorescence) and Hoechst (1 μg/mL, blue fluorescence). The images were taken at randomly selected areas to ensure the imaging conditions were consistent throughout the experiments. To ensure the reliability of the results, the experiments were repeated four times. Cells were imaged on an inverted laser scanning fluorescent microscope (Olympus) using a 60× oil immersion objective lens.

### The Cy5-PEG2 dye can effectively detect mitochondrial Morphology and Dynamics

Aβ_1-42_ peptide induces mitochondrial dysfunction and impacts mitochondrial morphology and dynamics. ^6^ Herein, it is crucial to determine whether the newly synthesized Cy5-PEG2 dye can effectively detect these changes. This requires additional experiments to evaluate the dye’s performance in cells treated with Aβ1-42 peptide and compare the results with control cells.

Primary neuronal cells were incubated with Aβ_1-42_ peptide at concentrations ranging from 0.1, 1, 10, and 100 μM for 24 hours. At concentrations of 100 μM, most of the mitochondria in primary neuronal cells were fragmented after being incubated with Aβ_1-42_ peptide for 24 hours. The cause of this fragmentation is likely due to the Aβ_1-42_ peptide-induced mitochondrial fission, which results in smaller organelles. This disintegration is likely due to the mitochondrial fission triggered by the high Aβ_1-42_ peptide concentration. The increased fission disrupts mitochondrial function, leading to cellular malfunction, and may play a role in neurodegenerative processes.^6^ However, an interesting phenomenon was observed in cells pretreated with Aβ_1-42_ peptide at a low concentration (0.1 μM). The mitochondria in low concentrations of Aβ_1-42_ peptide-pretreated cells appeared to have more tubular structures and were more interconnected (**Fig. 4**). This could be due to an adaptive response of cells triggered by low concentrations of Aβ_1-42_ peptide, which helps maintain mitochondrial dynamics and protect cell viability.

**Figure 4.**
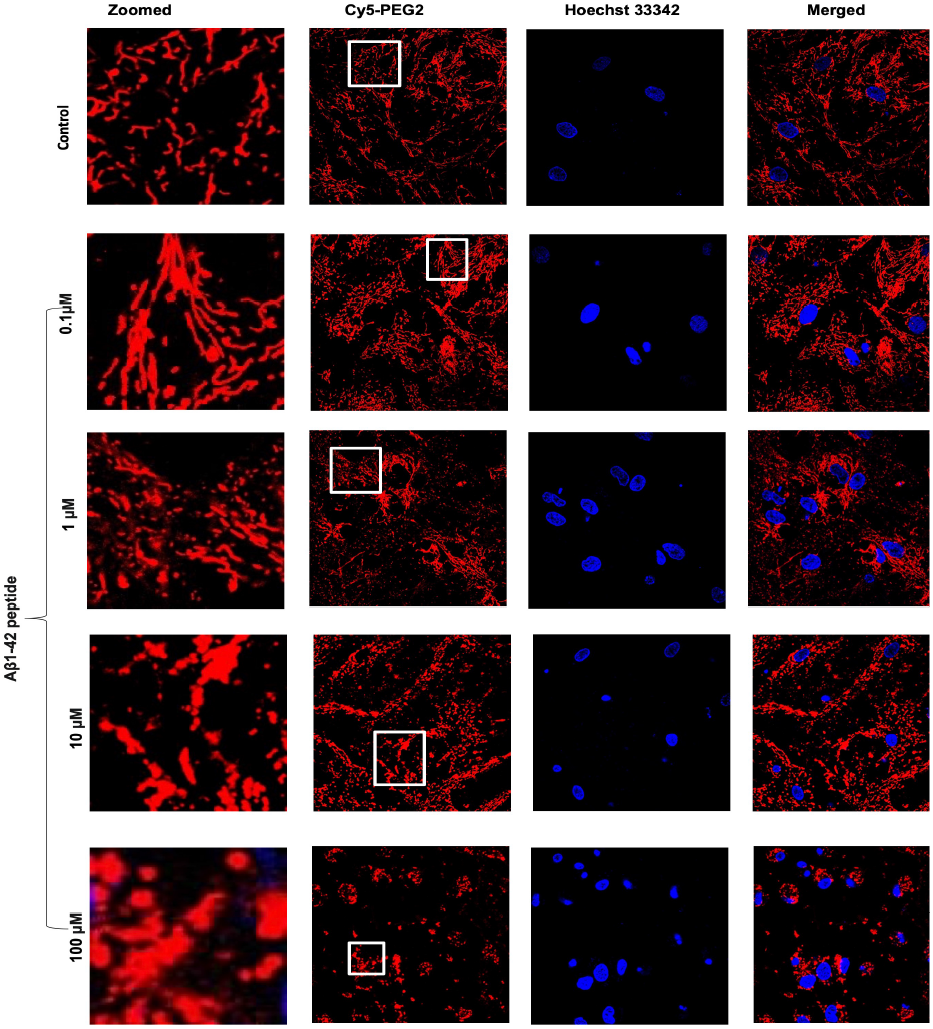
Representative confocal images of the primary neuronal cell lines in the presence of Aβ_1-42_ peptide at varying concentrations (0, 0.1, 1, 10, and 100 μM) for 24h. Cells were stained with Cy5-PEG2 (1 μM, red fluorescence) and co-stained with Hoechst (1 μg/mL, blue fluorescence). The images were taken at randomly selected areas to ensure the imaging conditions were consistent throughout the experiments. To ensure the reliability of the results, the experiments were repeated four times.

### The Cy5-PEG2 dye can effectively traverse the BBB and image brain cells’ mitochondria

To determine if the Cy5-PEG2 dye is suitable for brain imaging, it is necessary to test its ability to cross the BBB when administered intraperitoneal injection. The BBB is a selective barrier that prevents many therapeutic and imaging agents, including fluorescent dyes, from entering the brain. Administering the dye intraperitoneally allows us to evaluate its ability to reach the brain from the bloodstream. If the dye can cross the BBB and accumulate in brain tissue, it suggests that it has potential for use in brain imaging. We have confirmed that the Cy5-PEG2 dye can penetrate the BBB (**Fig. 5 and 6**).

**Figure 5.**
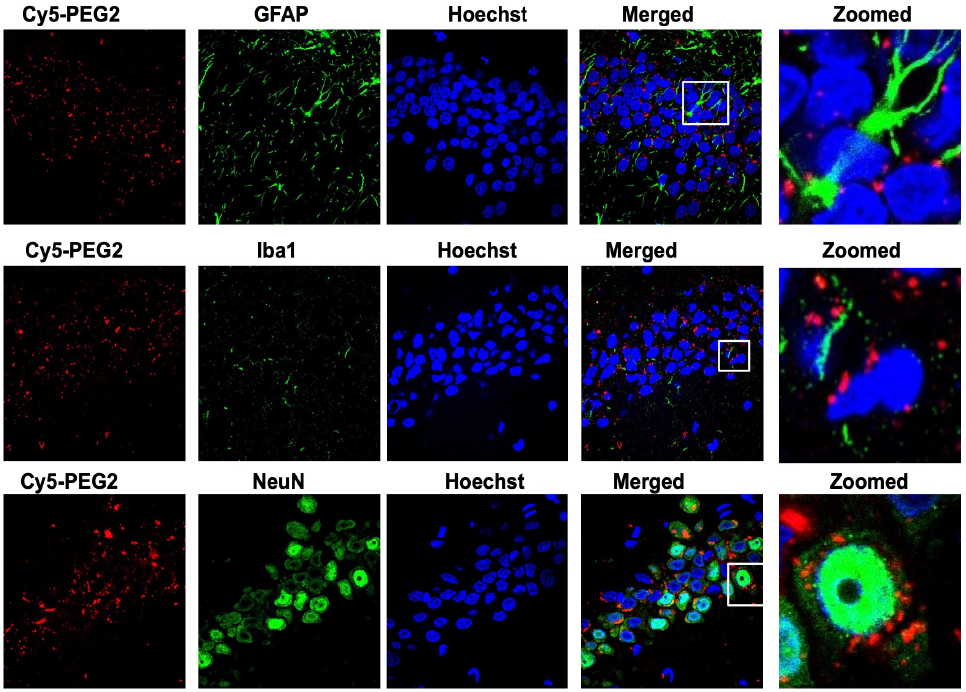
Visualization of the Cy5-PEG2 dye in the hippocampus regions: Representative confocal laser scanning images of rat brain sections fixed with 4% PFA. For the entire figure, the red channels represent the fluorescent dye, Cy5-PEG2; the green channel represents GFAP(row I), Iba1 (row II), and NeuN-positive (row III) cells in the brain sections, respectively. Zoomed-in pictures are shown on the right. The fluorescent images were obtained with a confocal laser scanning fluorescent microscope using a 60 × objective lens.

**Figure 6.**
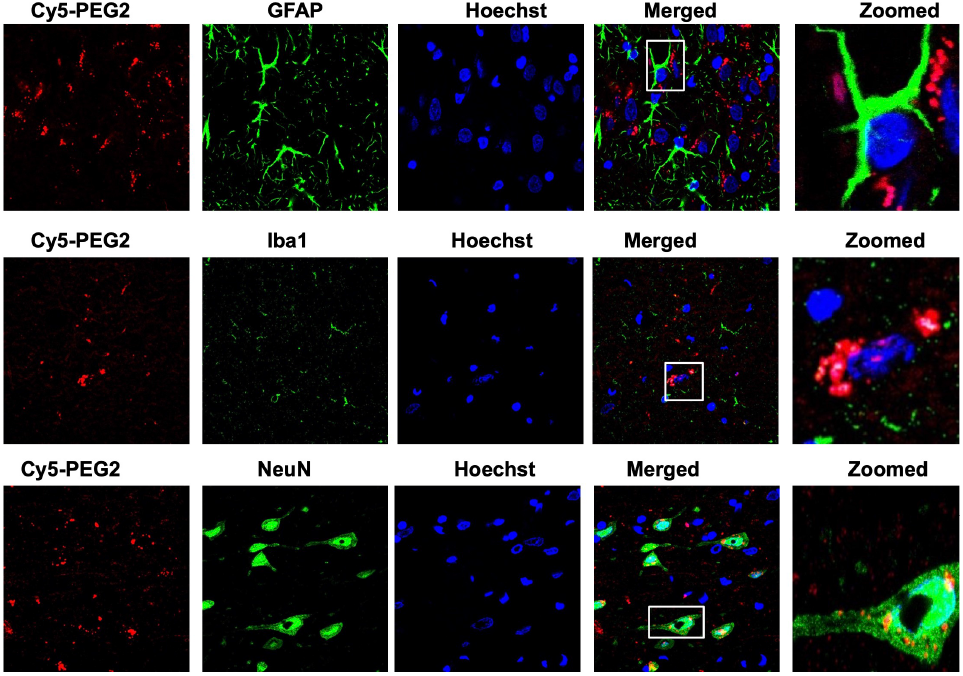
Visualization of the Cy5-PEG2 dye in the neocortex regions: Representative confocal laser scanning images of rat brain sections fixed with 4% PFA. For the entire figure, the red channels represent the fluorescent dye, Cy5-PEG2; the green channel represents GFAP(row I), Iba1 (row II), and NeuN-positive (row III) cells in the brain sections, respectively. Zoomed-in pictures are shown on the right. The fluorescent images were obtained with a confocal laser scanning fluorescent microscope using a 60 × objective lens.

As AD progresses, it affects various brain regions, leading to a decline in cognitive abilities and memory loss. The hippocampus is crucial for forming and consolidating memories, but the accumulation of Aβ peptide in this area can impair memory and the ability to create new memories.^9^ The neocortex, which includes the frontal, parietal, temporal, and occipital lobes, is also heavily affected by Aβ buildup, resulting in cognitive decline, language difficulties, and problems with executive functions. ^9^ In our present study, we observed the uptake and distribution of Cy5-PEG2 dye in these regions. We found that the red fluorescence of the Cy5-PEG2 dye was visible in both the hippocampus (**Fig. 5**) and neocortex (**Fig.6**), indicating that cells such as neurons and glial cells in these regions had absorbed this dye.

GFAP and IBA1 are biomarkers commonly used to assess the activation and response of glial cells in the central nervous system. Following brain injury or neurodegenerative conditions, GFAP expression increases. This upregulation is known as astrogliosis, a typical response to neuronal damage or inflammation in the brain.^10^ Monitoring GFAP levels can provide insights into the degree of glial activation and the extent of neural injury or damage caused by injected agents, such as Cy5-PEG2. Elevated GFAP expression may indicate a toxic response or neuroinflammation.

Iba1 is a protein only in microglia, the immune cells in the central nervous system. Microglia play an essential role in the brain’s immune response by detecting and monitoring pathogens, damaged neurons, and abnormal protein clusters. Microglia change their appearance and increase their Iba1 expression when activated, indicating their immune response and inflammation involvement. ^10^ Tracking Iba1 levels can evaluate microglial activation and neuroinflammation after injecting substances like Cy5-PEG2. An increase in Iba1 expression may suggest a potential toxic or inflammatory response.

By quantitating the levels of GFAP and Iba1 expression, we could assess the potential toxicity of the Cy5-PEG2 dye when administered intraperitoneal injection in SD rats. Our results indicate no significant rises in GFAP and Iba1 levels in the hippocampus and neocortex of Cy5-PEG2 injection rats compared to the rats that received vehicle control injection (data not shown), which suggests that the Cy5-PEG2 dye did not cause glial activation or inflammatory responses. Furthermore, there were no observable negative consequences or neurotoxicity on the central nervous system.

## Discussion

Live-cell imaging using fluorescent dyes like MitoTracker can help visualize mitochondria in various cell types. However, using these dyes for *in vivo* brain cell imaging is challenging due to specific limitations. For example, MitoTracker dyes need to penetrate plasma and mitochondrial membranes to label mitochondria, which is difficult in brain imaging due to the BBB, which affects their effectiveness in penetrating brain cells. Additionally, imaging deep brain regions can be challenging due to light scattering, absorption, and tissue autofluorescence, which can disrupt clear and specific labeling with MitoTracker dyes. Lastly, MitoTracker dyes can sometimes accumulate in other cellular compartments or structures, leading to potential false signals or confusion in interpretation. ^11^ To overcome these limitations, we aim to find new fluorescent dyes that target mitochondria and aid in studying these organelles, particularly for in vivo brain imaging.

Developing new fluorescent dyes that specifically target mitochondria in brain cells *in vivo* is a complex and challenging process. Brain cells have unique characteristics that pose specific obstacles, such as mitochondrial specificity, BBB penetration, photostability and brightness, biocompatibility, mitochondrial dynamics and morphology, long-term imaging and live-cell applications, and background fluorescence and autofluorescence.

Cyanine dyes are highly beneficial in bioimaging due to their distinct optical properties.^3,7^ They possess strong fluorescence, high quantum yields, and good photostability, making them ideal for various imaging techniques. By fluorescently labeling biomolecules like proteins, nucleic acids, and lipids, cyanine dyes allow for easy visualization and tracking of these molecules within living cells and tissues. In microscopy, researchers commonly use cyanine dyes as fluorescent probes to study cellular structures and dynamics with high resolution and sensitivity. Additionally, these dyes serve as optical imaging probes for specific applications, providing valuable insights into cellular processes and pathological conditions. Overall, cyanine dyes are excellent tools for bioimaging, with several practical applications in the field. Based on these positive results, we have created a range of new fluorescent dyes using cyanine for imaging mitochondria. This manuscript details the development, synthesis, and testing of one of these new fluorescent dyes, Cy5-PEG2, which can be used for both in vitro and in vivo imaging of brain cell mitochondria.

AD patients often have neurofibrillary tangles and senile plaque in their brains. Senile plaque is made up of Aβ peptides, while neurofibrillary tangles are comprised of hyperphosphorylated tau protein.^9^ The neocortex is typically the first area of the brain to display senile plaques, followed by the hippocampus and the rest of the brain in a centripetal pattern. The accumulation of Aβ peptides within the cells leads to this damage. Amyloid precursor protein (APP), a transmembrane protein expressed on chromosome 21 in neuron synapses, is critical for neuronal development. Aβ peptides are an abnormal proteolytic byproduct of APP. ^12^ In a healthy brain, APP undergoes alternative splicing, cleaving by α-secretase at or near lysine-16 in the Ab sequence on neuron surfaces.^12^ This releases the C-terminal site of soluble APP-alpha from the membrane and subsequent secretion, reducing Aβ protein deposition in cells. However, in the case of AD, APP is cleaved by βand γsecretase, leading to the formation of several isoforms of Aβ peptides ranging in length from 39 to 43 residues. ^12^ Aβ fibrils found in senile plaques are usually the Aβ species that end at amino acid 42 (Aβ42). These fibrils are longer and more hydrophobic than other Aβ species, making them more prone to aggregation and toxicity.

Previous studies suggested that accumulating Aβ peptides in the brain can have severe consequences.^12,13^ It can trigger the production of reactive oxygen species (ROS), initiate cell death processes, and increase neurotoxicity. Aβ peptides have been identified as the cause of dendritic degeneration and loss of synapses, which ultimately leads to neuronal death in the rat hippocampus.^12^ Additionally, in vitro studies using cells from PC12 cells (rat pheochromocytoma ^14,15^ and SH-SY5Y cells ^16,17^ (human neuroblastoma) have demonstrated that extracellular fibrillar aggregates of Aβ peptides are neurotoxic. However, these cells are unsuitable for neurodegenerative disease research due to their instability and inability to create a diverse population of neurons.

Our present study utilized primary neuronal cells derived directly from the nervous system to understand how Aβ peptide affects mitochondrial morphology and dynamics. These cells are a biologically relevant model that closely mimics the brain’s cellular environment, making it easier to study the molecular mechanisms underlying AD. To evaluate this effect, we used the Cy5-PEG2 dye to monitor the alterations in the mitochondrial morphology and function of primary neuron cells.

Mitochondria are essential organelles known as the “powerhouses” of the cell because they produce energy through oxidative phosphorylation, which generates adenosine triphosphate (ATP). They also play a crucial role in other cellular processes, such as apoptosis, calcium homeostasis, and cellular signaling. Mitochondria are dynamic organelles that can undergo fusion and fission processes, contributing to maintaining mitochondrial function and overall cellular health.

Our present study examined the effects of the Aβ_1-42_ peptide on mitochondrial morphology and dynamics. The Aβ_142_ peptide can increase fragmented mitochondria at high concentrations. This process may result from increased cellular stress or damage. Mitochondria fission can lead to mitochondria fragmentation, impairing their function and ultimately contributing to cellular dysfunction. Interestingly, the Aβ_1-42_ peptide was observed to increase the percentage of elongated mitochondria at low concentrations (Supporting Information). Mitochondrial fusion might be an adaptive response to cellular stress, possibly aimed at improving mitochondrial function by sharing resources and repairing damaged mitochondria.

AD is characterized by the accumulation of amyloid beta peptides in the brain, forming plaques and subsequent neurodegeneration. Interestingly, depending on concentration, the Aβ_1-42_ peptide can affect mitochondrial morphology and dynamics. One significant finding is that high concentrations of the Aβ_1-42_ peptide can cause mitochondria fission, which is the splitting of mitochondria into smaller units. Fission can lead to various cellular stress responses, energy imbalance, and cell death when fission is not regulated correctly. This is particularly relevant to AD, as disrupted mitochondrial dynamics can contribute to neuronal dysfunction and degeneration.^13^ On the other hand, low concentrations of the Aβ_1-42_ peptide can induce mitochondria fusion, which is the merging of mitochondria. This process helps maintain mitochondrial function and cellular health by redistributing mitochondrial components, sharing mitochondrial DNA, and rescuing damaged mitochondria.

The findings in our present study suggest that modulating mitochondrial dynamics could be a potential strategy to counteract the adverse effects of the Aβ peptide. By promoting fusion or inhibiting excessive fusion, we may be able to mitigate the impact of high Aβ_1-42_ concentrations or prevent the accumulation of dysfunctional mitochondria. It is important to note that the effects of Aβ_1-42_ on mitochondria are concentration-dependent. This highlights the complex and often biphasic nature of cellular responses to pathological factors. Therefore, studying dose-response relationships is crucial when investigating the impact of such molecules on cellular processes. These findings also underscore how the Aβ_1-42_ peptide can induce cellular stress responses contributing to disease progression. Dysregulation of mitochondrial dynamics is not limited to AD but is a common theme in various neurodegenerative disorders, including Parkinson’s and Huntington’s.^2,13^ These observations provide valuable insights into the intricate relationship between amyloid beta peptide and mitochondrial dynamics. Further research in this area may lead to potential therapeutic strategies to counteract the detrimental effects of amyloid beta peptide on mitochondrial function and neuronal health.

Our investigation into the toxicity profile of Cy5-PEG2 dye in the brain involved carefully examining its impact on glial activation, neuronal health, and potential neuroinflammation. After conducting our study, we did not identify any significantly increased levels of GFAP and Iba1, indicating that the injected Cy5-PEG2 did not trigger glial activation or inflammatory responses. Moving forward, assessing changes in neuronal morphology, synaptic integrity, and cell viability would be beneficial to gain a more comprehensive understanding of this dye’s toxicity profile.

## Conclusion

A new fluorescent dye, Cy5-PEG2, has been developed to accumulate in mitochondria, enabling in vivo brain imaging selectively. This advancement is significant as it can penetrate the blood-brain barrier (BBB), making it an excellent tool for monitoring mitochondrial dynamics in living cells. It is expected to be used to study mitochondrial dynamics and function in the context of whole-brain physiology and disease progression.

## Limitations

Based on the data we have collected thus far, the Cy5PEG2 dye we administered does not significantly impact the expression levels of GFAP and Iba1 in the brain tissues. This suggests that the dye does not activate glial cells or cause neuroinflammation. Furthermore, we did not detect any toxicity in the markers we analyzed. However, it may be beneficial to investigate other markers of inflammation or neurotoxicity to achieve a more comprehensive evaluation of this dye.

## AUTHOR INFORMATION

### Funding Sources

This project is patricianly supported by AHA 1807047 (Bi), NIHR15HL150703 (Shan), NIHR01HL163159 (Shan).

## Notes

### Competing Interest Statement

The authors have declared no competing interest.

